# Zero-preserving imputation of scRNA-seq data using low-rank approximation

**DOI:** 10.1101/397588

**Authors:** George C. Linderman, Jun Zhao, Yuval Kluger

**Affiliations:** Program in Applied Mathematics, New Haven, CT 06511; Computational Biology and Bioinformatics, New Haven, CT 06511; Department of Pathology Yale University, New Haven, CT 06511

## Abstract

Single cell RNA-sequencing (scRNA-seq) methods have revolutionized the study of gene expression but are plagued by dropout events, a phenomenon where genes actually expressed in a given cell are incorrectly measured as unexpressed. We present a method based on low-rank approximation which successfully replaces these dropouts (zero expression levels of unobserved expressed genes) by nonzero values, while preserving biologically non-expressed genes (true biological zeros) at zero expression levels. We validate our approach and compare it to two state-of-the-art methods. We show that it recovers true expression of marker genes while preserving biological zeros, increases separation of known cell types and improves correlation of simulated cells to their true profiles. Furthermore, our method is dramatically more scalable, allowing practitioners to quickly and easily recover expression of even the largest scRNA-seq datasets.

## 1. INTRODUCTION

Measurement of gene expression in individual cells requires amplification of truly minute quantities of mRNA, resulting in a phenomenon called “dropout,” in which an expressed transcript is not detected and hence assigned a zero expression value. Dropout results in an excess of zeros, some of which are biological zeros (i.e. truly not expressed), whereas others are technical zeros (i.e. expressed but not measured). The profound sparsity of scRNA-seq data can be detrimental to downstream analyses, so several methods have been developed to address it [1–7]. However, the most popular of these methods do not distinguish between technical and biological zeros, treating all zeros as missing data, and resulting in a matrix where biological zeros are often also completed. Furthermore, these methods do not scale to the size of many scRNA-seq datasets, where hundreds of thousands to millions of cells are analyzed.

We present Adaptively-thresholded Low-Rank Approximation (ALRA): a highly scalable method for recovery of true scRNA-seq expression levels, which takes advantage of the non-negativity and correlation structure of expression matrices to selectively impute technical zeros. Our approach (Figure 1) rests on a key observation about low-rank approximation. First, we assume that the underlying true expression matrix is non-negative, low-rank, and contains many zeros, but where all entries associated with dropouts are non-zero. The result of a scRNA-seq experiment is then an even sparser matrix, since many values are incorrectly measured as zero due to the dropout effect. We compute a low-rank approximation of this matrix using the singular vector decomposition (SVD) and observe that the values corresponding to zero in the original matrix are symmetrically distributed around zero. Due to the symmetry, the negative values provide an estimate for the error distribution of the elements corresponding to true zeros. As such, we set to zero all the elements of each row with absolute value smaller than the most negative value on that row, ensuring that the only completed values are those are above the maximal expression level of the error distribution of the biologically non-expressing genes.

**Figure 1.**
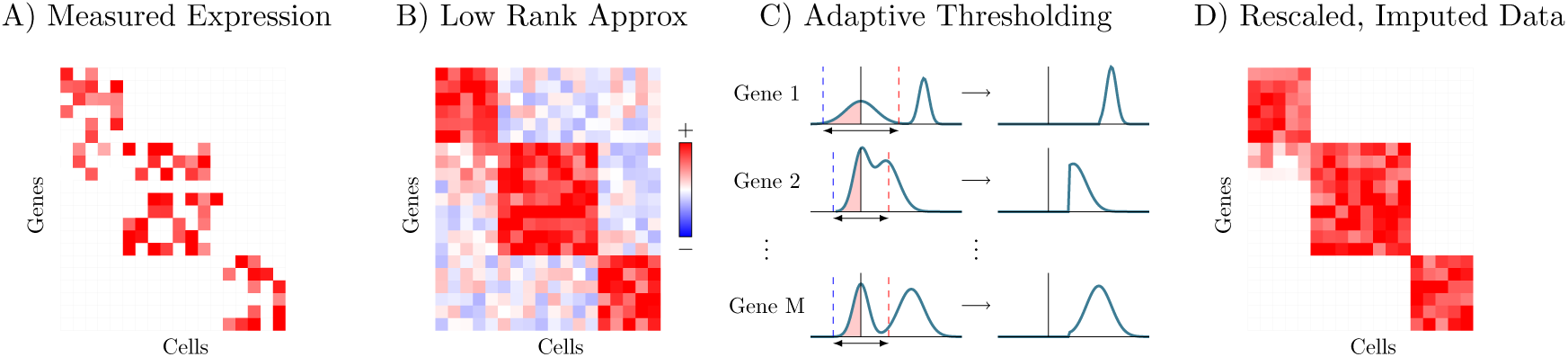
Overview of the ALRA imputation scheme. Measured expression matrix (A) contains technical zeros (in each block) and biological zeros (outside each block). Low rank approximation via SVD results in a matrix (B) where elements corresponding to biological zeros for each gene are symmetrically distributed around 0, so that thresholding (red dotted line) by the absolute value of the most negative value (blue dotted line) of each gene (C) restores the biological zeros (D).

We emphasize the simplicity of our approach: it is SVD followed by a thresholding scheme that takes advantage of the non-negativity of the true matrix. Remarkably, this simple approach outperforms other more complex methods for recovery of scRNA-seq expression data, as we demonstrate below. We compare ALRA to a diffusion-based approach called MAGIC [2], which exhibited the best relative performance across datasets among methods for imputation of scRNA-seq data in a recent review [6]. We also compare to SAVER, a recently published approach which assumes a statistical model for read counts and estimates parameters using LASSO regression [1].

## 2. RESULTS

### 2.1. ALRA preserves biological zeros in simulated and real datasets

We assessed ALRA’s ability to selectively complete technical zeros using scRNA-seq of 10 purified populations of peripheral blood monocytes (PBMCs) generated by Zheng et al. (2017) [8]. We ran ALRA, SAVER, and MAGIC on the merged matrix, and focused our attention on B cells, natural killer cells, cytotoxic T cells, and helper T cells. These four cell types are well characterized and there are known marker genes specific to each. For example, CR2 (CD21), NCAM1 (CD56), CD8A, and CD4 are specific to B cells, NK cells, cytotoxic T cells, and helper T cells, respectively. Using these genes, we demonstrate SAVER’s ability to preserve biological zeros.

After low rank approximation (the first step of ALRA), every element in the matrix is non-zero. However, the elements of the matrix that correspond to biological zeros (e.g. the gene CR2 in NK cells, cytotoxic T cells, or helper T cells) are symmetrically distributed around zero. By thresholding at the red dotted lines, which are symmetrical mirror image of the most negative value of each gene (blue dotted lines), we ensure that the biological zeros are preserved. For CR2 and NCAM1, this process preserves the biological zeros while completing all of the technical zeros. However, for CD4 and CD8A, the distributions of elements corresponding to biological zeros and technical zeros overlap, and hence some of the elements corresponding to technical zeros are also set to zero (Figure 2A). For this reason we consider ALRA to be a “conservative” imputation method of scRNA-seq data: it does not impute biological zeros, even at the cost of missing some technical zeros.

**Figure 2.**
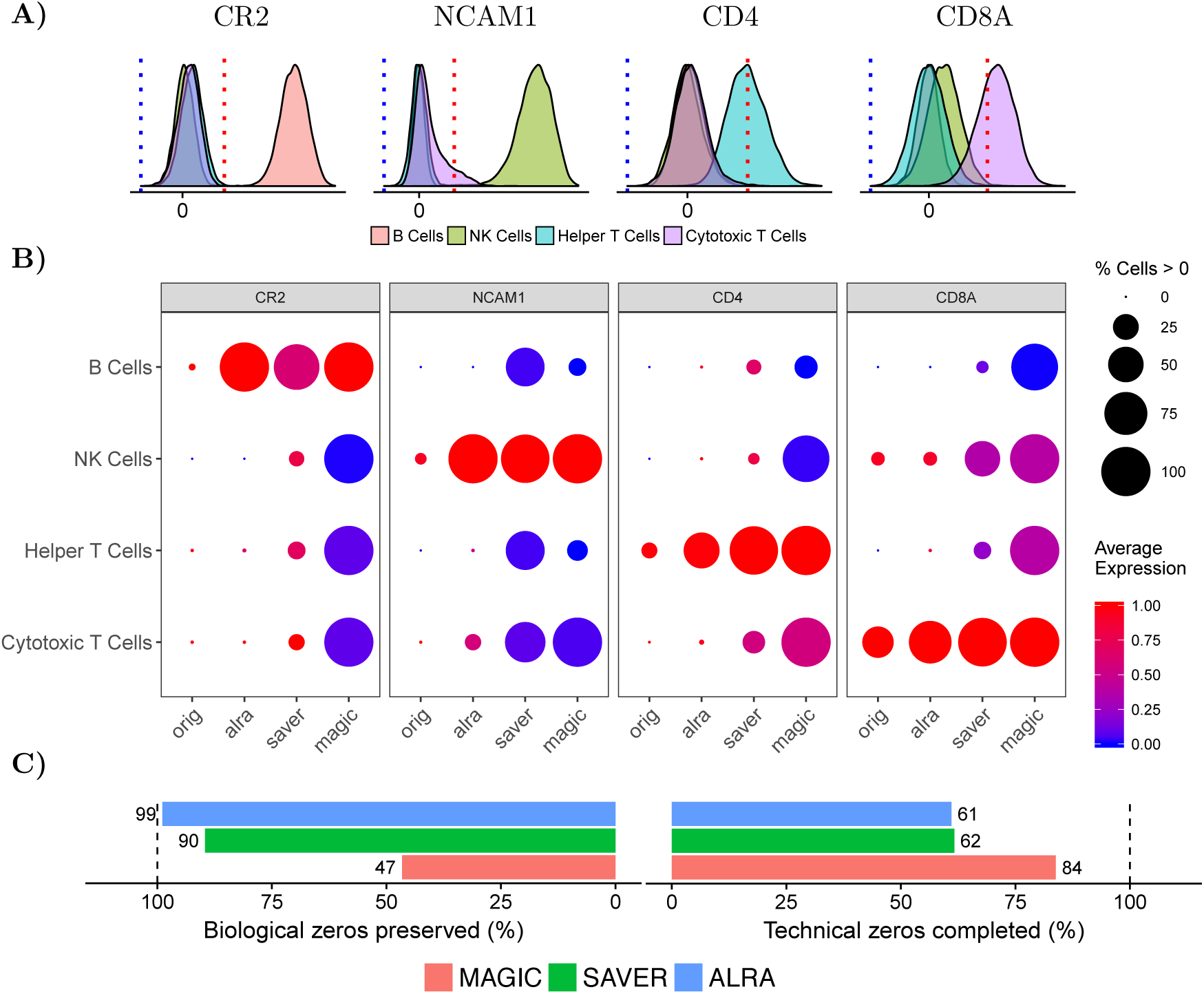
Preserving biological zeros when completing missing values in purified PBMC populations. A) After low-rank approximation of purified PBMC populations, ALRA recovers biological zeros of each gene by thresholding the expression values of that gene in all cells at the absolute value of the most negative expression level. B) ALRA preserves biological zeros in the purified PBMC dataset while completing a comparable number of technical zeros to other methods. Expression values are normalized to the max of each gene. C) Percentage of biological zeros preserved and technical zeros completed in a simulated scRNA-seq dataset.

As shown in Figure 2B, ALRA recovers the true sparsity pattern of known marker genes in these four purified peripheral blood monocyte populations. We note that some cytotoxic T cells with NK cell markers (i.e. CD8+CD56+ cells) are present in the peripheral blood, which justifies completion of NCAM1 in a small fraction cytotoxic T cells [9]. In contrast, SAVER and MAGIC successfully complete the technical zeros, but also complete many zeros expected to be biological, based on the known cell type specificity of these markers

To assess ALRA’s ability to selectively complete technical zeros (while preserving biological zeros), we used bulk RNA-seq of 9 immune populations from the ImmGen consortium [10] to simulate scRNA-seq expression data. For each population, we generated single cell profiles by sampling counts from a multinomial distribution (as in [11,12]) with probabilities proportional to expression in the bulk profile. The resulting matrix (see Methods) of 17,580 genes and 5000 cells consisted of 93% zeros, and each zero can be identified as a technical zero (gene actually expressed in corresponding bulk RNA-seq sample) or biological zero (gene actually not expressed in the corresponding bulk RNA-seq sample).

As shown in Figure 2, ALRA preserves 99% of the true biological zeros while completing a large proportion of the non-zeros. In contrast, MAGIC completes all of the biological zeros, with about 47% positive values. SAVER fares better, but still incorrectly completes more biological zeros than ALRA. We evaluated the overall performance of these approaches using the balanced accuracy, obtained by averaging the biological zeros preservation rate and the technical zeros completion rate. For ALRA, MAGIC, and SAVER, the balanced accuracy are 80%, 65%, and 75% respectively.

### 2.2. ALRA improves separation of cell types

We demonstrate the effect of ALRA on the visualization of scRNA-seq data using t-SNE on a dataset containing 65,539 mouse visual cortex cells from Hrvatin et al. (2018) [13], each of which had been classified by the authors into 30 distinct cell types. Figure 3 compares the t-SNE before and after ALRA, which demonstrates that distinct cell types. Figure 3 compares the t-SNE before and after ALRA, which demonstrates that several cell types can be more clearly separated. To quantitatively confirm this result, we trained a supervised random forest to classify the cells into cell types, and we compare the out-of-bag error before and after ALRA. The error is more than halved from 6.3% error in the original data to 2.6% error after ALRA, confirming that ALRA improves the separation of the cell types.

**Figure 3.**
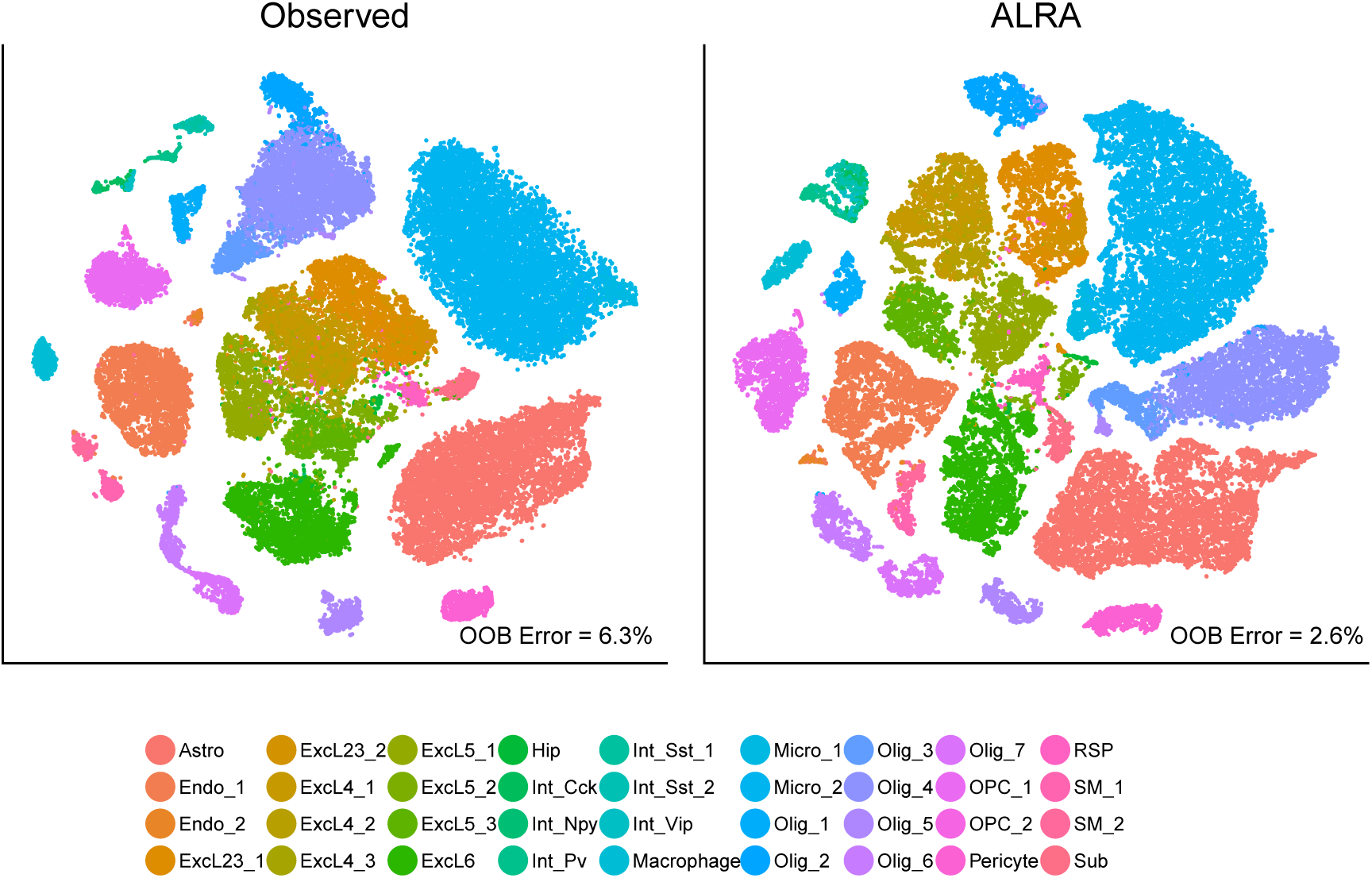
ALRA improves separation of previously annotated cell types in mouse cortical cells as shown by comparing t-SNE before (left) and after (right) imputation by ALRA.

In Huang et al. (2018), SAVER was also shown to clearly improve separation between cell types in a random sample of 7,387 cells from the same dataset. We could not compare our result for the full dataset of 65,539 cells in Figure 3 to SAVER’s, because the latter estimated that on a single-core it would take many days to run on the full dataset. For completeness, we run ALRA on the same subsample as SAVER and show that both improve separation of cell types as compared to the original data (Figure S2). We note that Huang et al. found SAVER to improve the t-SNE of all 4 tested datasets as compared to MAGIC, hence we omit direct comparison to MAGIC here.

### 2.3 ALRA recovers true expression distribution

Using the same simulated dataset as before, we computed the correlation between each cell’s expression profile and the bulk RNA-seq profiles for the 9 immune cell populations they were generated from. If the correlation between a cell and its corresponding bulk RNA-seq sample is larger than the others, we say it is correctly classified. We would expect that after recovery, the correlation between a cell and its corresponding bulk RNA-seq sample would be clearly larger than its correlation to the other samples, and hence the classification rate would improve.

As shown in Figure 4A, most cells are classified correctly in the original data, but just barely. ALRA successfully increases the correlation of each cell with its corresponding bulk RNA-seq sample more than with the other cell type bulk profiles. MAGIC and SAVER, however, do not improve the correlations; in fact, the misclassification error is higher after recovery using these methods.

**Figure 4.**
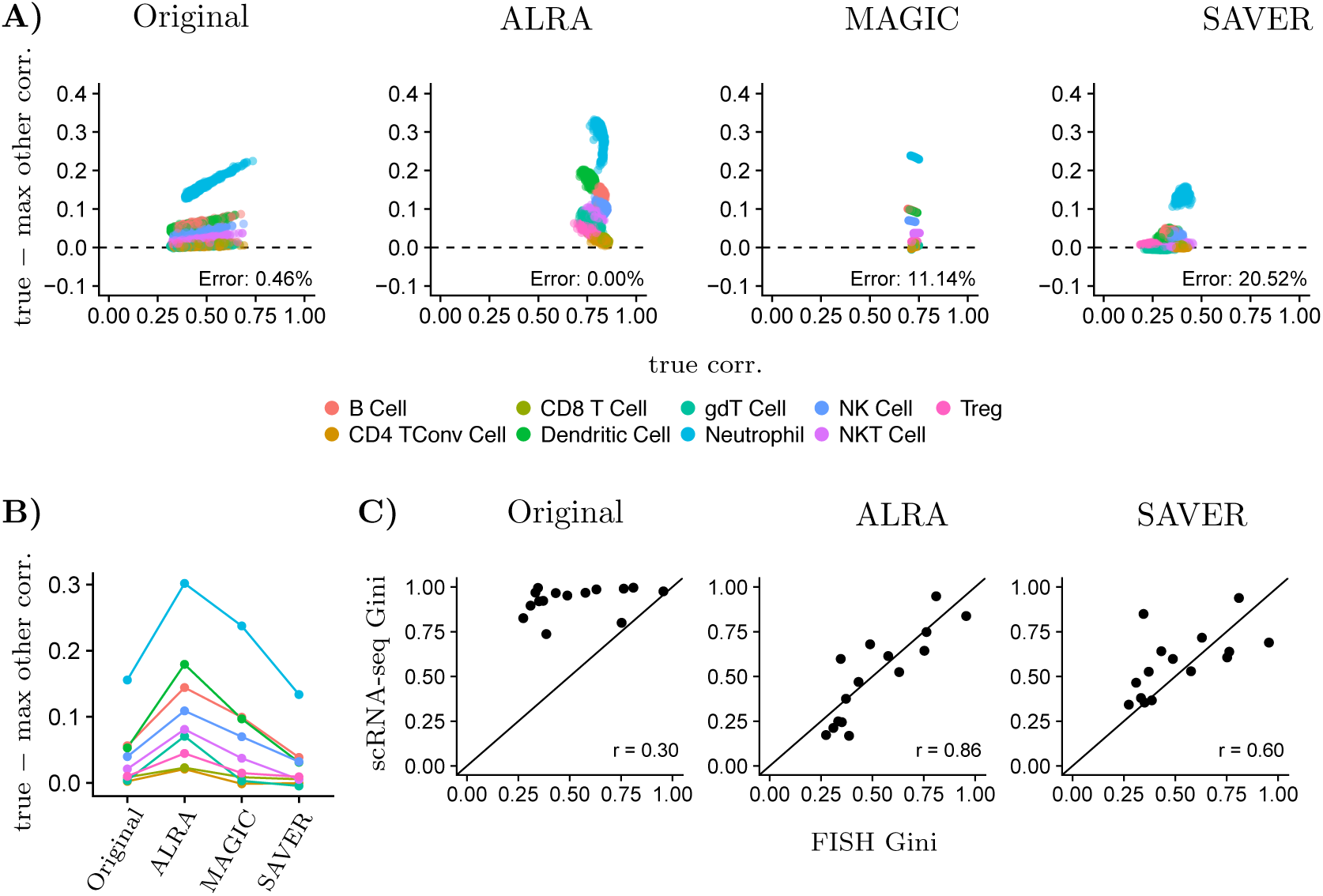
Validating recovered expression profiles. A) Correlation between simulated immune cell and its true profile from bulk RNA-seq (“true correlation,” x-axis) vs. difference between true correlation and max correlation of simulated immune cell with other profiles from bulk RNA-seq (y-axis). Points where lying below zero are considered misclassified. B) Y axis in (A) averaged over cells for each population. C) Gini coefficient for 15 genes computed from scRNA-seq and FISH measurements of the same melanoma cell line.

As in Huang et al., we further evaluated our approach by comparing ALRA’s completion of Drop-seq data to RNA florescence in situ hybridization (FISH) of the same population of cells. In Torre et al. (2018), 8,498 cells from a melanoma cell line were sequenced by Drop-seq [14]. RNA FISH was also used to measure the expression of 26 markers in cells of the same cell line. Although the gene expression profiles of individual cells cannot be compared across these two datasets, the distribution of gene expression should be consistent, because the cells were sampled from the same cell line.

Huang et al. (2018) applied SAVER to the Drop-seq data and demonstrated that, as compared to the raw data, the estimated distribution of expression values is more consistent with the FISH distributions [1]. Their analysis computes the Gini coefficient of each gene, a measure of gene expression variability, which should be consistent across the Drop-seq and FISH technologies. Specifically, they showed that the Gini coefficients of a subset of marker genes in the FISH dataset were more consistent (correlation 0.6) with the SAVER estimates than either the raw Drop-seq data (correlation 0.3) or other completion methods. Our approach out-performs SAVER on this metric, obtaining a correlation of 0.86. We note that Huang et al. (2018) already found their method to perform better than MAGIC in this regard, and hence we omit direct comparison here.

Furthermore, SAVER’s performance depends on several filtering steps, such that only 15 of the 26 genes were considered in the analysis. The marker genes were also each normalized by GAPDH expression, in order to control for technical biases between technologies. We demonstrate that ALRA’s performance is not sensitive to these steps (Figure S3) and exhibits a correlation of 0.82 on the full dataset. These results suggest that ALRA more accurately recovers the true distribution of the data and that its performance is less sensitive to preprocessing.

### 2.4. ALRA is highly scalable

ALRA computes a rank-*k* approximation to the expression matrix using randomized SVD, and hence it can scale to arbitrarily large datasets. In Figure 5A we compare the runtime of ALRA, MAGIC, and SAVER on subsets of mouse visual cortex cells from Section §2.2. The resulting matrices ranged from 1,000 to 50,000 cells and each had 19, 155 genes. ALRA dramatically out-performs the other methods, taking *∼*40 minutes on 50,000 cells. In contrast, MAGIC requires over 4 hours from the same number of cells. SAVER required over 12 hours to run on 5,000 cells, and hence was not attempted on larger subsets.

**Figure 5.**
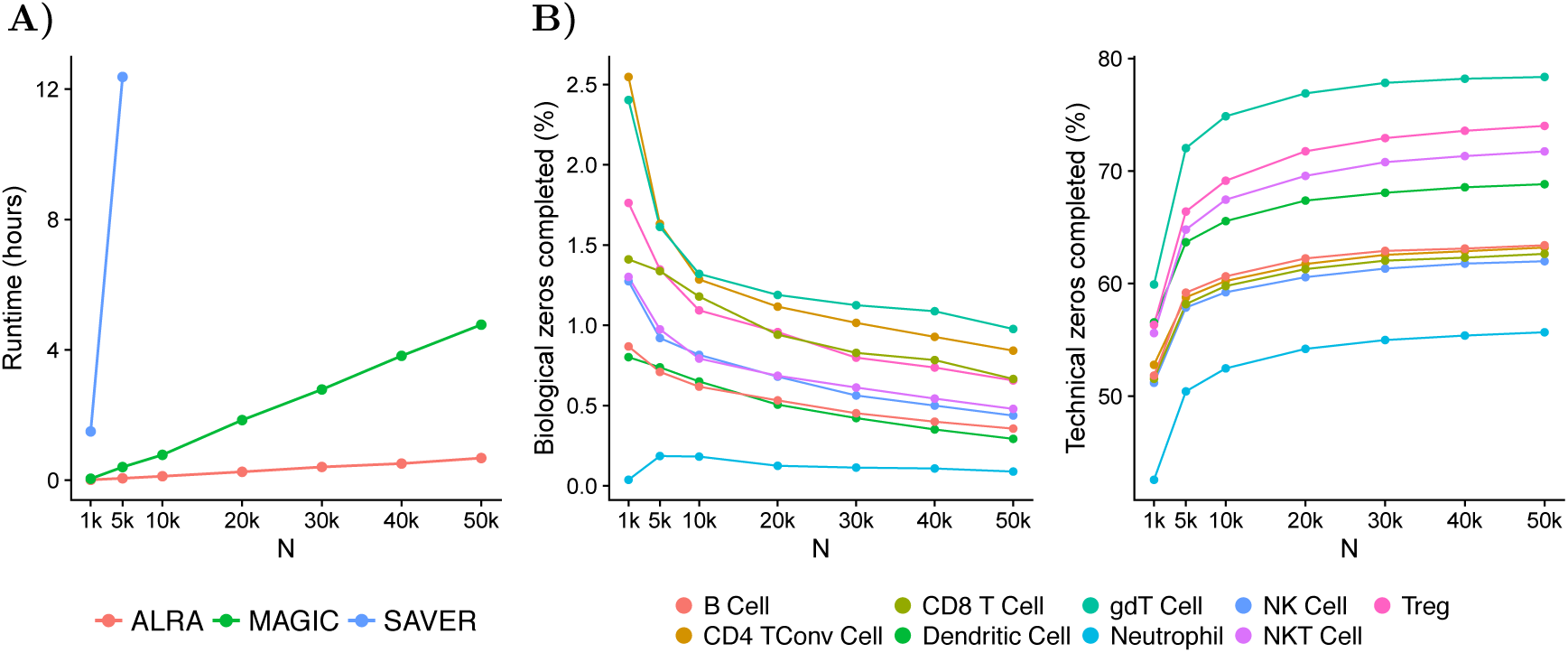
Runtime and performance of ALRA as number of cells increases. A) Runtime of ALRA is compared to MAGIC and SAVER using expression matrices of *N* cells and ∼19, 000 genes. B) ALRA’s performance improves as the number of cells increases, as demonstrated using a simulated scRNA-seq dataset where biological and technical zeros are known.

ALRA’s scalability is more than convenience; its speed obviates the need for subsampling prior to recovery of expression. Using the simulated scRNA-seq dataset from before, we demonstrate that ALRA’s results improve when applied to increasingly large datasets. In particular, we show that the number of biological zeros incorrectly completed in each simulated cell population decreases as *N* increases, while the number of technical zeros correctly completed increases (Figure 5B).

## 3. DISCUSSION

We present ALRA, a method for recovery of true gene expression from dropped out scRNA-seq data, which preserves true biological zeros while substantially improving downstream analyses through imputation of technical zeros. In particular, we showed that ALRA improves separation of cell types in both t-SNE and the original high dimensional space, improves correlation of simulated cells to their true profiles, recovers true expression of marker genes while preserving biological zeros, and scales dramatically better than other methods. We also note that ALRA has only one parameter, the approximate rank *k* of the matrix, which is selected automatically based on statistics of the spacings between consecutive singular values.

The key assumption underlying ALRA is that the true scRNA-seq data matrix is low-rank, meaning that expression values lie on a linear subspace of lower dimension (as compared to the dimensions of the matrix). It has long been appreciated [15–18] that genes do not act independently, but rather in concert, forming groups of highly correlated genes often referred to as gene modules. Therefore, the true expression matrix modelled as a low-rank matrix. It is because of this low rank structure that nearly every scRNA-seq pipeline first reduces dimensionality to the top principal components (e.g.[19]): the variation of the signal in the expression data can be captured by a relatively small number of principal components.

Although the true expression matrix may be low-rank, the measured expression matrix is only approximately low-rank. That is, the elements are close to lying on a lower dimensional subspace, but not exactly. The optimal rank-*k* approximation to this matrix can be easily found using the singular value decomposition (SVD)[20], resulting in a matrix where every element is non-zero. This simple process is sometimes referred to as SVD imputation, and it was applied to scRNA-seq as a baseline method in [1].

Our method is motivated by a simple observation: the non-zero values incorrectly assigned to true zeros using SVD imputation are symmetrically distributed around zero. Therefore, any element larger in magnitude than the most negative value should not correspond to a biological zero. Conversely, values smaller in magnitude than the most negative value could correspond to biological or technical zeros, and hence we restore them to their original value. The result is a completed matrix where the biological zeros are preserved and where every completed value can be trusted to be actually non-zero.

This crucial observation is of independent theoretical interest. Let *M* be a rank-*k*, non-negative matrix and let Ω be the set of indices (*i, j*) such that *M*_*i,j*_ = 0. Suppose we observe a matrix *X* where some values, with indices Θ, are set to zero. That is, *X*_*i,j*_ = 0 if (*i, j*) ∈ Θ and *X*_*i,j*_ = *M*_*i,j*_ otherwise. Now, letting 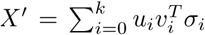 be the truncated SVD of *X*, and letting *i*_0_ be any row index, we hypothesize that the distribution of 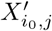 for (*i*_0_, *j*) ∈ Ω ∩ Θ is symmetric around zero. That is, that the error distribution of values corresponding to 0 in any row of the low-rank approximation of a perturbed non-negative, low-rank matrix is symmetric around zero.

Low-rank approximation induces the simplest kind of low-rank matrix completion (LRMC) and is closely related to a rich theory that has developed for completion of missing values in low-rank matrices (e.g. [21–23]). For instance, a LRMC method based on minimization of the nuclear norm [24] was shown to be effective for the imputation of dropped out values in scRNA-seq expression matrices [6]. However, direct application of LRMC algorithms to scRNA-seq data treats all the zeros as missing and completes them all, just like the low-rank approximation step used in ALRA. After completion using LRMC methods (e.g. minimization of the nuclear norm), it is not clear if the error distribution of the elements corresponding to biological zeros is symmetric around zero (as in low rank approximation). If so, then an adaptive thresholding scheme similar to ours could be applied as a post-processing step to other LRMC methods.

An R implementation of ALRA is freely available at https://github.com/KlugerLab/ALRA, and the codes for all analyses in this paper are available at https://github.com/KlugerLab/ ALRA-paper.

## 4. ACKNOWLEDGEMENTS

The authors would like to thank Stefan Steinerberger and Ofir Lindenbaum for many useful discussions.

G.C.L. acknowledges support by the NIH NHGRI Grant #F30HG010102. Y.K. acknowledges support by NIH grant #1R01HG008383-01A1.

## 5. METHODS

### 5.1. ALRA procedure

Given an expression matrix, we first perform library normalization to 10,000 UMIs per cell and then take a log to obtain the normalized expression matrix. The ALRA procedure consists of three steps: First, we use the R package rsvd (version 0.9) [25] to compute the near-optimal rank-*k* approximation of the normalized expression matrix, setting the parameter for number of additional power iterations *q* = 10. Next, we threshold each gene of the resulting matrix by the absolute value of that gene’s most negative entry. Finally, we rescale the resulting values such that the mean and standard deviation of the non-zero values of each gene in the resulting matrix match that of the original matrix. A small number of originally non-zero values are below the threshold and get set to zero in this process, so we restore them to their original values.

The choice of *k* is critical, and we provide a heuristic which was used to choose *k* for all experiments. Noise typically manifests as a long tail of singular values, and the goal is to choose *k* such that all singular values *σ*_1_, …, *σ*_*k*_ correspond to signal, whereas the rest of the values are noise. We first use rsvd to compute the first 100 singular values, with the default setting of *q* = 2 additional power iterations. We compute the spacings between consecutive singular values, denoted as *s*_*i*_ = *σ*_*i*_ – *σ*_*i*–1_ from *i* = 2, …, 100. As can be seen in the plots of *s*_*i*_ for two different datasets (Figure 6), we aim to choose the largest *k* where *s*_*k*_ is significantly different from spacings in the tail. Specifically, we compute the mean *µ* and standard deviation *σ* of *s*_80_, …, *s*_100_, and then for each difference *s*_*i*_ we compute the probability *p*_*i*_ of observing a value as extreme as *s*_*i*_ from a normal distribution with mean *µ* and standard deviation *σ*. We then let *k* be the largest index such that *p*_*i*_ < 1 ×10^*-*10^. That is, *σ*_*k*_ is the smallest singular value such that *σ*_*k*+1_ –*σ*_*k*_ is significantly different than the spacings between singular values in the tail. We emphasize that this is a heuristic, albeit one that appears to be quite effective; we note that other methods for estimating the optimal rank of truncated SVD have been explored in the statistical literature (e.g. [26–28]).

**Figure 6.**
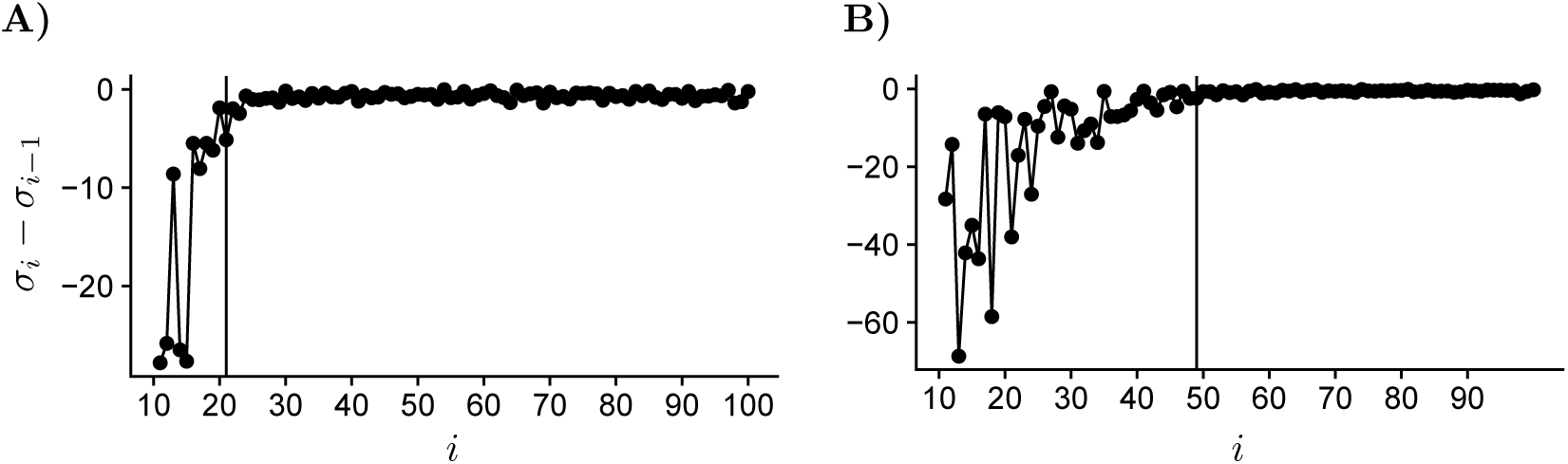
Differences between consecutive singular values. A) the rank was set to *k* = 20 in the purified PBMCs experiment. B) Rank was set to *k* = 48 for the mouse visual cortex cells experiment.

### 5.2. Purified PBMC analysis

Filtered expression matrices were obtained for FACS purified B cells, CD14 monocytes, CD34+ cells, CD4 helper T cells, regulatory T cells, naive T cells, memory T cells, CD56 natural killer cells, cytotoxic T cells, and naive cytotoxic T cells from the 10X Genomics website. The matrices were merged, resulting in an expression matrix with 94,655 cells and 32,738 genes. The matrix was filtered to include cells expressing more than 400 genes and genes expressed in more than 100 cells, resulting in a matrix with 83,992 cells and 12,776 genes. We applied SAVER to this count matrix (as SAVER takes the raw data and performs its own normalization), and sampled from the estimated posterior distribution using the function ‘sample.saver’. We ran ALRA and MAGIC on the library and log normalized the expression matrix. The *k* parameter for ALRA was chosen to be *k* = 20 using the procedure described in Section §5.1.

### 5.3. Simulated scRNA-seq data

To evaluate ALRA’s performance, we generated simulated scRNA-seq data from ImmGen [10] deep-sequenced bulk RNA-seq data of nine spleen immune cell types (B, CD4 TConv, CD8 T, Dendritic, gtD, neutrophil, NK, NKT, and Treg cells). Raw reads of bulk RNA-seq data from ImmGen [10] were mapped to the mm10 genome with STAR [29] and annotated by HTSeq [30] to generate counts per gene.

The simulation is based on a multinomial model, as in [11,12]. Enumerating the bulk RNA-seq samples as *i* = 1, …, 9 and genes as *j* = 1, …, *m*, let 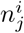 denote the read counts of gene *j* in the *i*th sample. Normalizing, we parameterize the multinomial distribution with an *m*-dimensional probability vector *p*^*i*^, where 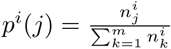.

To generate the gene expression profile for the *𝓁*th cell, we choose one of the 9 cell types with equal probability and denote the cell type of the *𝓁*th cell by *c*_*𝓁*_ ∈ {1, …, 9}. The associated probability vector is then *p*^*c𝓁*^. In order to obtain a realistic distribution of read counts, we randomly sample a cell from the purified PBMC dataset, and let *N*_*𝓁*_ be the read count in that cell. Finally, we sample an *m*-dimensional vector of counts from the multinomial distribution parameterized by *N*_*𝓁*_ and *p*^*c𝓁*^. The process is then repeated for as many cells as specified by the simulation.

This simulation data enables us to distinguish true biological zeros and technical zeros based on whether or not the gene is actually expressed in the corresponding bulk RNA-seq sample.

### 5.4. Improved separation of mouse cortical cells

The scRNA-seq expression matrix of mouse visual cortex cells from [13] was obtained and filtered as in Huang et al., so as to compare with SAVER’s result. Specifically, genes with mean expression less than 0.00003 and non-zero expression in less than 4 cells were excluded, resulting in an expression matrix with 19,155 genes and 65,539 cells. ALRA was run with *k* = 48 (as chosen by procedure described in §5.1) on the subset of 48,244 cells that were classified into cell types in the original study. Variable genes were identified for both the observed and ALRA-completed data using Seurat [31]. The fast t-SNE implementation of [32] (FIt-SNE) was then used to compute the embedding of the top 48 principal components of the observed and ALRA-completed data.

To quantitatively compare the separability of cell types, we trained a random forest to classify each cell as its cell type, as we would expect better separated cell types to be easier to classify. We used the R package ranger [33] with default settings and compared the out-of-bag error before and after expression recovery using ALRA.

### 5.5. FISH experiments

We obtained the Drop-seq dataset from [14], consisting of 8,640 cells and 32,287 genes. The RNA FISH measurements from the same paper were also obtained, consisting of 26 genes across 7,000 to 88,000 cells (depending on the gene). We then computed the Gini coefficient of each gene in RNA FISH, original Drop-seq, Drop-seq after SAVER, and Drop-seq after ALRA (Figure S3).

In addition, to directly compare with analysis published by the authors of SAVER, we followed their same filtering procedure, removing cells with library size greater than 20,000 or less than 500, and also removing genes with mean expression less than 0.1. The resulting dataset contained 8,498 cells and 12,241 genes, with 16 genes shared between Drop-seq and FISH. As in Huang et al., cells in the bottom and top tenth percentiles of the housekeeping gene GAPDH were filtered out, and all genes were then normalized by GAPDH expression. We then computed the Gini coefficient of each of the other 15 genes (excluding GAPDH) in RNA FISH, original Drop-seq, Drop-seq after SAVER, and Drop-seq after ALRA, using this filtered subset (Figure 4B).

### 5.6. Runtime comparison

We subsetted the filtered Hrvatin et al. data into random subsets of sizes 5,000, 10,000, 20,000, 30,000, 40,000 and 50,000, and compared the runtime of MAGIC, SAVER and ALRA on each subset. All experiments were performed on a single core, with default parameters. Notably, SAVER’s option of ‘do.fast’ was set to true. For ALRA, the normalization, initial computation of 100 singular vectors with *q* = 2 using rsvd to choose *k*, and the actual completion procedure were all included in the timings. We note that randomized SVD, the most time-consuming step of ALRA, can be easily parallelized (e.g. as in [34]) and lead to further performance gains.

## 6. SUPPLEMENTAL FIGURES

**Figure S1.**
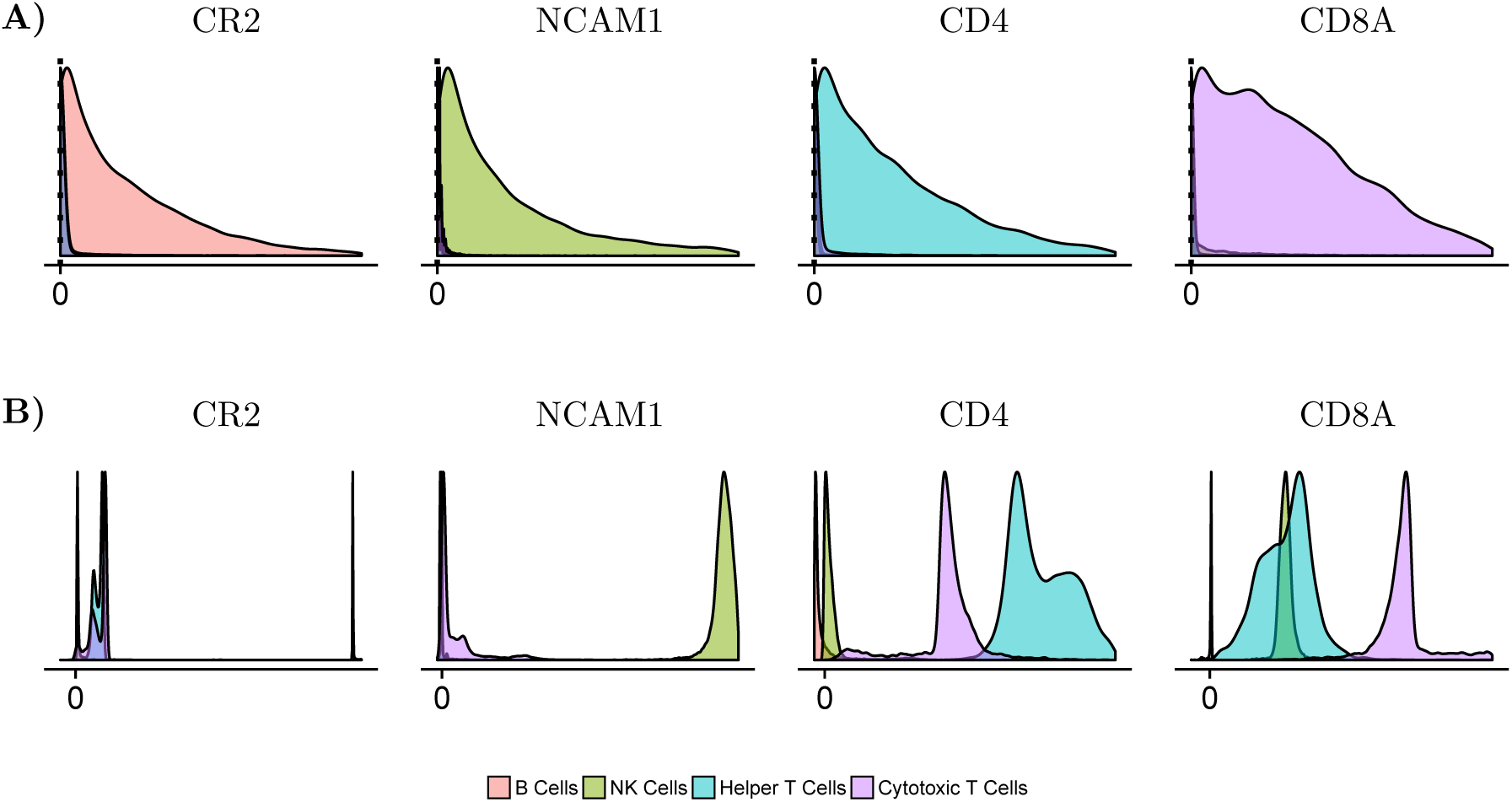
Imputation of purified PBMC dataset (Most extreme 1% are excluded for plotting purposes). A) SAVER’s output B) MAGIC’s output.

**Figure S2.**
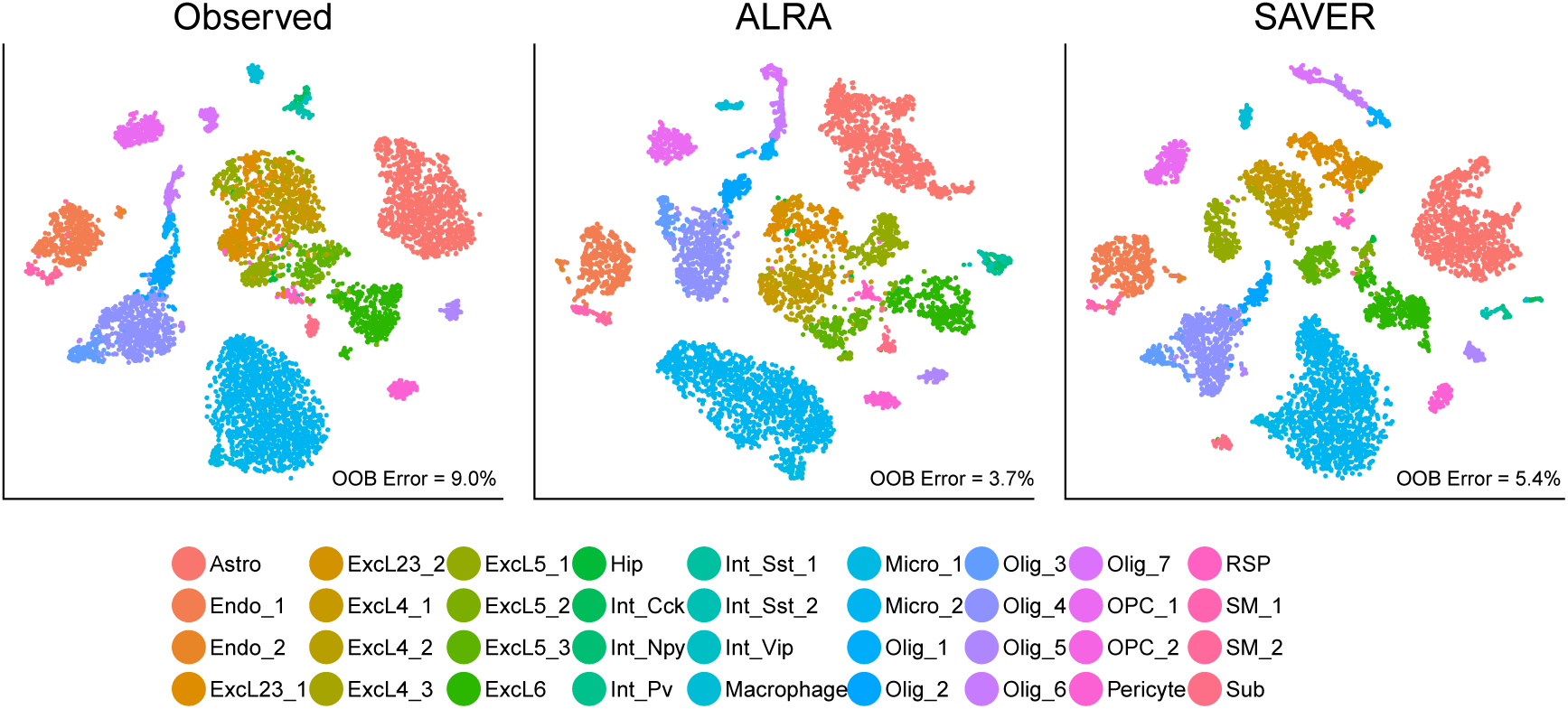
ALRA and SAVER both improve separation of previously annotated cell types on the same subset of mouse cortical cells as analyzed in Huang et al. (2018).

**Figure S3.**
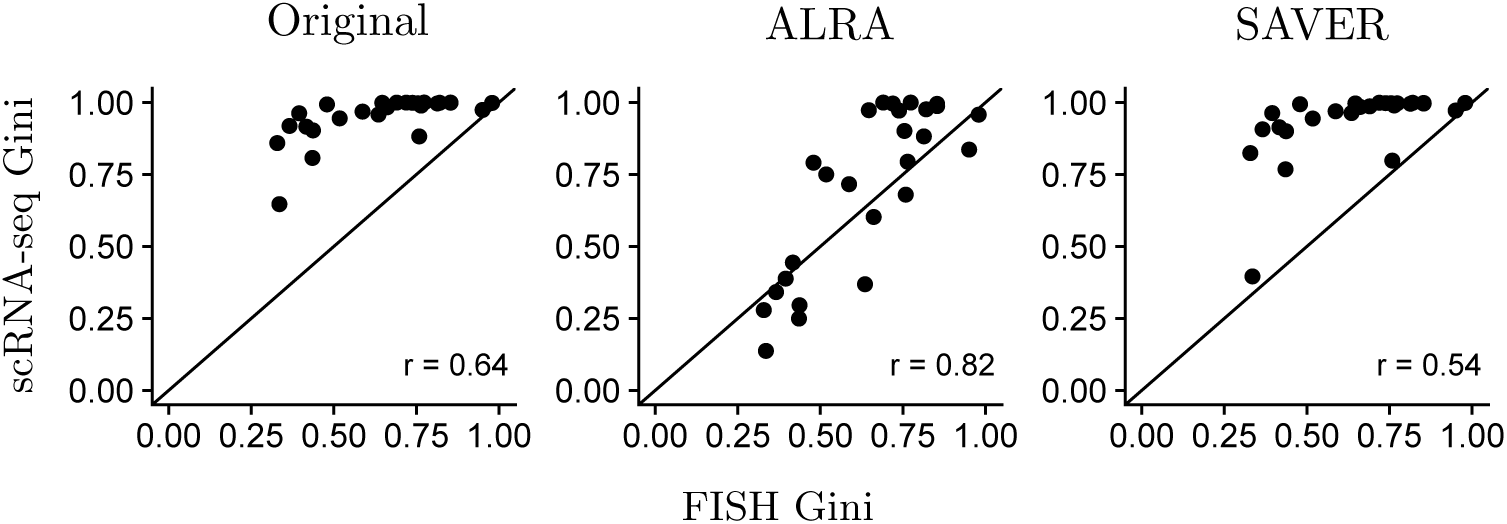
Gini coefficient for all 26 genes computed from scRNA-seq and FISH measurements of the same melanoma cell line.

